# Structural Mechanism of TRPC3 Channel Activation by the Moonwalker Mutation

**DOI:** 10.64898/2026.04.03.716262

**Authors:** Jiahe Zang, Yongmei Tan, Yikun Chen, Wenjun Guo, Xiaole Zhao, Hailin Peng, Lei Chen

**Affiliations:** State Key Laboratory of Membrane Biology, College of Future Technology, Institute of Molecular Medicine, Peking University, Beijing Key Laboratory of Cardiometabolic Molecular Medicine, Beijing 100871, China; Academy for Advanced Interdisciplinary Studies, Peking University, Beijing 100871, China; Center for Nanochemistry, Beijing Science and Engineering Center for Nanocarbons, Beijing National Laboratory for Molecular Sciences, College of Chemistry and Molecular Engineering, Peking University, Beijing 100871, China; Peking-Tsinghua Center for Life Sciences, Peking University, Beijing 100871, China; National Biomedical Imaging Center, Peking University, Beijing, 100871, China

## Abstract

TRPC3 is a calcium-permeable, non-selective cation channel that is activated by DAG. It is expressed in several tissues, especially in the cerebellum, and has been implicated in various human diseases. Despite recent progress in understanding the structural mechanism of TRPC3, how the channel opens remains elusive. Here, we present structures of hTRPC3 in an agonist-free resting state, determined using a DAG-binding site mutant. We also present the structure of hTRPC3 in a DAG-bound open state, determined using a constitutively active “moonwalker” (T561A) mutant. These structures, together with electrophysiological results, reveal that the T561A mutation activates hTRPC3 by disrupting a polar interaction with N652. A newly formed π-bulge in S6 leads to rotation and outward tilting of the lower half of S6, resulting in dilation of the pore and thus channel opening. Agonist DAG stabilizes hTRPC3 in the open conformation. BTDM exerts its inhibitory effect by pushing S5 and S6 back to the center to close the pore, while preserving the π-bulge. These results shed light on the opening mechanism of hTRPC3.

## Introduction

Transient Receptor Potential Canonical (TRPC) proteins are a family of receptor-activated, non-selective cation channels that play crucial roles in cellular signaling ^1^. Upon activation, these channels mediate the influx of sodium and calcium ions, leading to membrane depolarization and the initiation of diverse calcium-dependent signaling cascades ^1^. Within this family, TRPC3, TRPC6, and TRPC7 form a subfamily characterized by high sequence homology and a shared mode of activation, being directly activated by the second messenger diacylglycerol (DAG) ^2,3^. These channels are implicated in a wide array of essential physiological processes, ranging from neuronal development and synaptic transmission to cardiovascular regulation and glomerular filtration ^1^. Consequently, they have emerged as promising therapeutic targets for conditions such as cardiac hypertrophy, neurological disorders, and cancer ^4^.

TRPC3, in particular, exhibits a distinct expression pattern, with notably high levels in the cerebellum, where it is critical for the proper development and function of Purkinje cells ^5^. The physiological importance of TRPC3 is underscored by the identification of a gain-of-function mutation, T561A, which results in impaired dendritic arborization of Purkinje cells and leads to cerebellar ataxia, manifesting as a distinctive moonwalker gait abnormality in mice ^5^. In addition to its function in Purkinje cells, TRPC3 has been increasingly recognized for its involvement in immune responses, cancer progression, and various cardiovascular and neurological disorders, highlighting its potential as a therapeutic target ^6,7^.

Significant progress has been made in understanding the structural basis of TRPC3 function. Early structural studies on the homomeric TRPC3 channel elucidated its overall architecture, revealing a tetrameric assembly composed of a transmembrane domain (TMD) and a large cytosolic domain (CTD) ^8–10^. Subsequent high-resolution structures identified key regulatory elements, including an inhibitory calcium-binding site (CBS) within the CTD that mediates negative calcium feedback (CBS1), and an activating calcium-binding site (CBS3) located in the voltage-sensor-like domain (VSLD) of the TMD ^11^. Most recently, the binding modes of several TRPC3 agonists, such as DAG, the synthetic small molecule GSK1702934A, and the endogenous modulator 4n, have been characterized, providing molecular insights into how agonists interact with the channel ^12^.

Despite these advances, a critical gap in our understanding remains: all available structures of TRPC3, and closely related TRPC6, including those with agonists bound, have captured the channel in a closed, non-conducting state. Therefore, the precise structural rearrangements that underlie the opening of the TRPC3 channel remain enigmatic. Here we present the structure of hTRPC3 in the agonist-free resting state and the DAG-bound open state of the constitutively active moonwalker (T561A) mutant, thereby elucidating the opening mechanism of hTRPC3.

## Results

### The structure of hTRPC3 in the agonist-free resting state

The DAG analogue oleoyl-2-acetyl-sn-glycerol (OAG) activates the hTRPC3 channel with a time course similar to typical ligand-gated ion channels, displaying slow but distinctive activation and desensitization phases (Fig. 1a). Although previously determined structures of TRPC3 all contain agonists at the DAG-binding L2 site located at the subunit interface ^8,9,11,12^, these structures are in a non-conductive, closed state. It is possible that these structures represent a pre-open state, in which the agonist is bound but the channel has not yet transitioned to the open state, or a desensitized state, in which the pore has closed following agonist-elicited opening. Therefore, we sought to determine the structure of human TRPC3 (hTRPC3) in the absence of agonist to capture its resting state structure.

**Figure 1.**
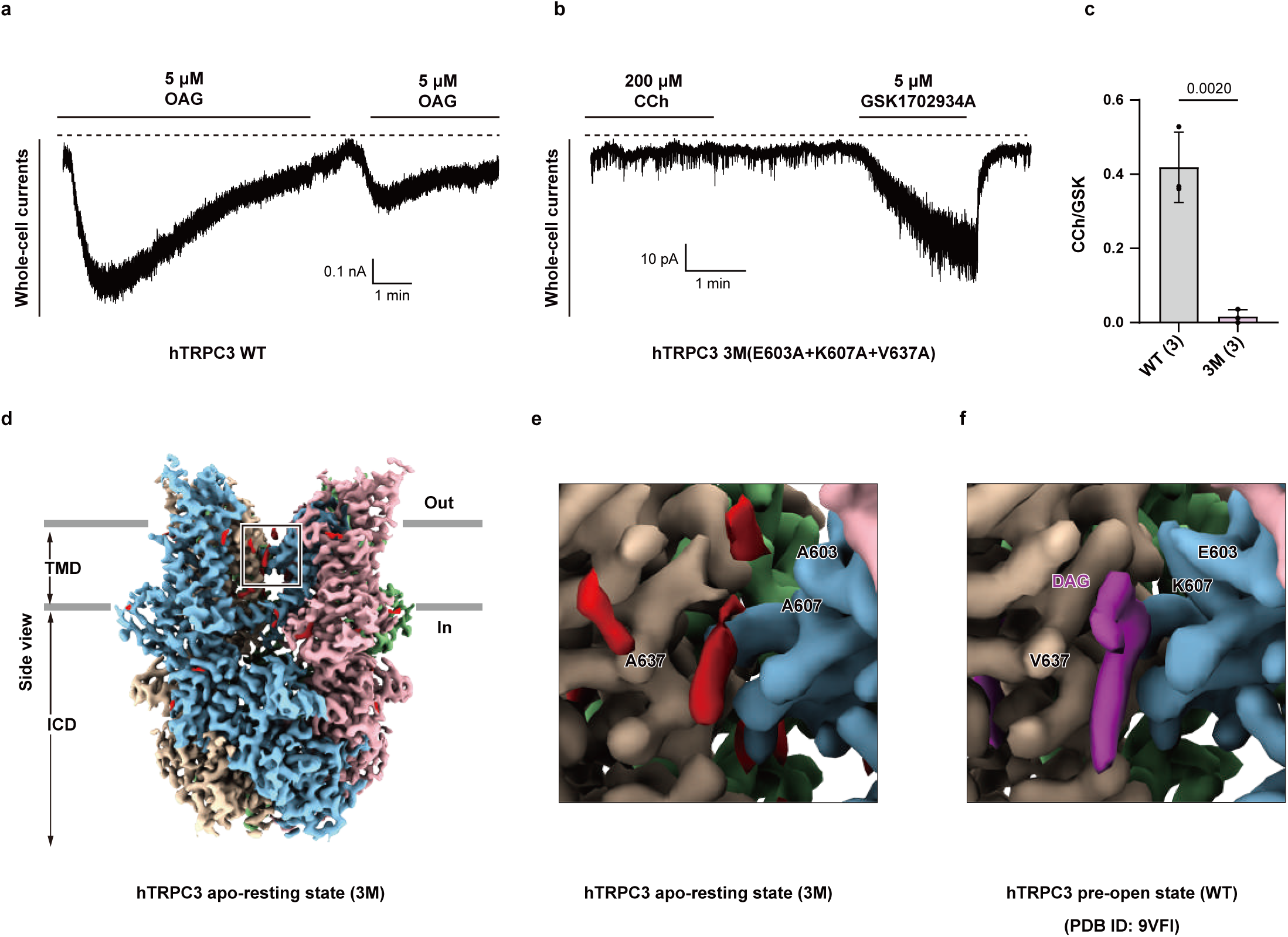
The structure of the hTRPC3 3M mutant in the DAG-free resting state. a, Whole-cell currents of the wild-type hTRPC3 recorded at −60 mV. The dashed line denotes the position of zero current. The recording has been repeated three times with similar results. b, Whole-cell currents of the wild-type hTRPC3 recorded at −60 mV. The dashed line denotes the position of zero current. The recording has been repeated three times with similar results. c, The ratio of whole cell peak currents activated by CCh and GSK1702934A in wild-type and 3M (E603A+K608A+V637A) mutant recorded at −60 mV. The data are shown as the mean ± SD. The number of replicates in each group is indicated in parentheses. The data were analyzed using t-tests with p-values indicated above the corresponding groups. d, Cryo-EM density map of the hTRPC3 3M mutant shown in side view. Subunits A, B, C, and D are colored in blue, pink, green, and yellow, respectively. Unmodelled residual densities are shown in red. Box denotes the L2 site, where E603A, K608A, and V637A mutations are located. e, Close-up view of the electron density of hTRPC3 3M mutant at the L2 DAG binding site boxed in panel d. f, Close-up view of the electron density of wild-type hTRPC3 (EMD-65025) at the L2 DAG binding site. DAG is colored in purple.

To achieve this, we combined three mutations, E603A, V637A, and K607A, which we previously identified to be at the DAG-binding pocket and important for DAG-binding ^12^ to generate a construct, hTRPC3-3M. This construct can be activated by synthetic activator GSK1702934A but showed no activation by mAChRs agonist carbachol (CCh) (Fig. 1b-c), confirming that its DAG-binding ability is greatly impaired. To minimize the artifact of detergent solubilization and retain the natively bound lipids, we purified the hTRPC3-3M protein using 4F-peptide solubilization and subsequent nanodisc assembly ^13^. We determined its structure to a resolution of 2.8 Å (Fig. 1d, Supplementary Fig. 1, and Supplementary Table 1). As expected, no DAG density was observed at the L2 site (Fig. 1e); instead, the DAG-binding site was occupied by an unknown lipid with distinct density morphology (Fig. 1e-f), suggesting this structure represents the apo-resting state of hTRPC3. Importantly, this structure is overall similar to the DAG-bound wild-type hTRPC3 channel purified in a similar way (PDB ID: 9VFI) ^13^, supporting the hypothesis that all of the previously determined agonist-bound structures ^12^ represent a pre-open state rather than the desensitized state.

### The structure of the hTRPC3 T561A mutant in the open state

We hypothesized that the open probability of wild-type hTRPC3 in the presence of agonists, such as DAG, 4n ^14^, or GSK1702934A ^15^ is not high enough for us to capture its open state^12^. Therefore, we sought to use a gain-of-function mutant to shift the conformational equilibrium toward the open state. Previous studies have identified the T561A (moonwalker) mutation, which renders mouse TRPC3 more active ^5^. We found that the T561A mutant is constitutively active and retains sensitivity to the high-affinity inhibitor BTDM ^9^, but not SAR7334 ^16^(Fig. 2a). To minimize the cytotoxicity of the constitutive activity of hTRPC3 T561A during cell culture, we expressed the T561A mutant in the presence of BTDM, which was then extensively washed away during protein purification. hTRPC3 T561A protein was extracted from the cell membrane using 4F peptide to preserve natively bound lipids ^13^. The purified T561A mutant was reconstituted into nanodiscs ^13^ and subjected to single-particle cryo-EM analysis (Supplementary Fig. 2-3 and Supplementary Table 1).

**Figure 2.**
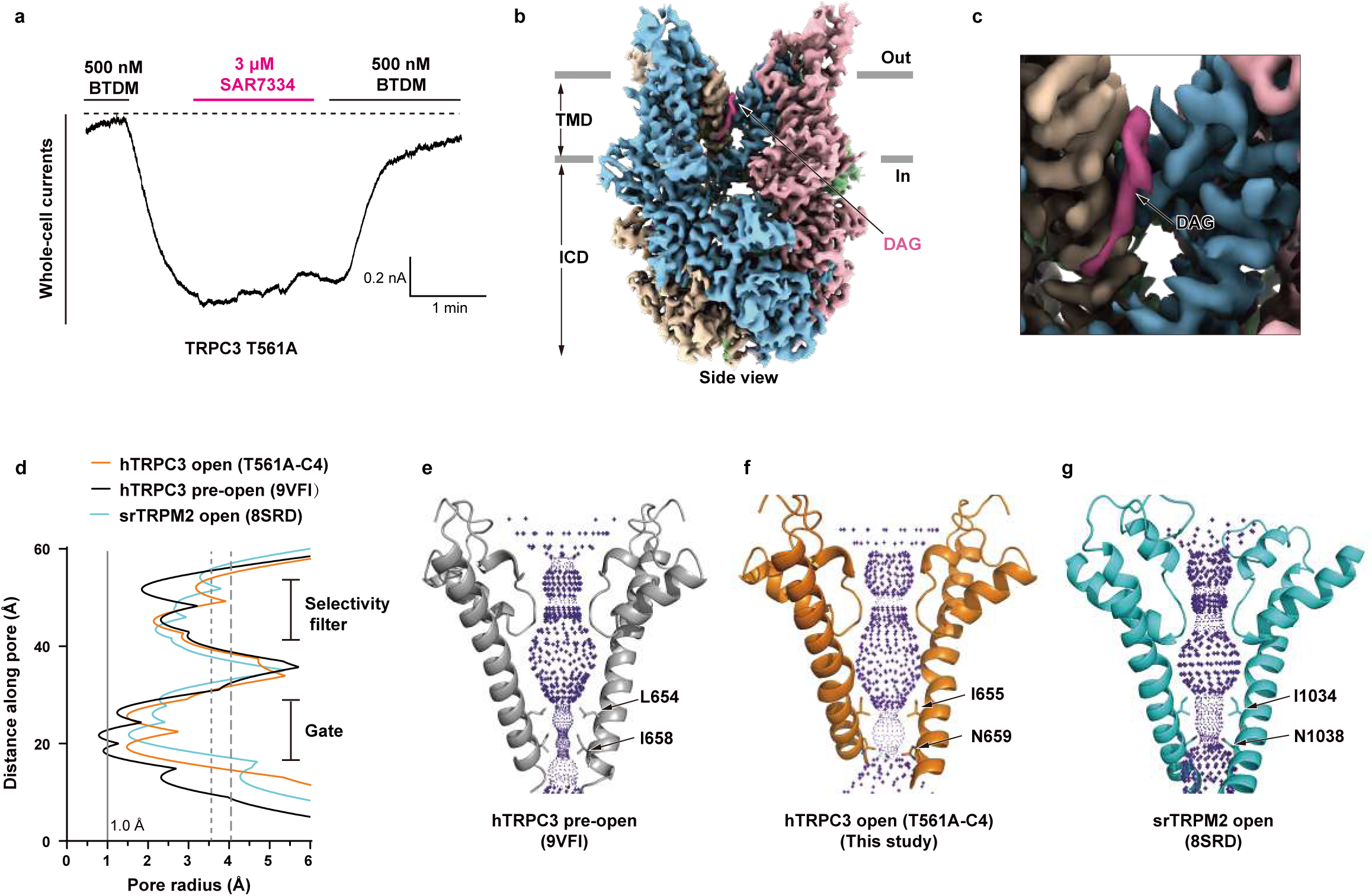
The structure of hTRPC3 T561A mutant in the open state. a, Whole-cell currents of the hTRPC3 T561A mutant recorded at −10 mV. The dashed line denotes the position of zero current. The recording has been repeated three times with similar results. b, Cryo-EM density map of the hTRPC3 T561A mutant (T561A-C4) shown in side view. Subunits A, B, C, and D are colored the same as Fig. 1d. DAG is colored in purple. c, Close-up view of panel b at the L2 DAG binding site. d, Pore radius calculated using HOLE2. The gray solid line denotes the radius of the dehydrated calcium ion (1.0 Å). The gray dashed lines denote the radius of the hydrated Na^+^ (dotted) and Ca^2+^ ions (dashed). e, Pore profile of hTRPC3 in the pre-open (PDB ID: 9VFI) calculated using HOLE2. Only the S6 of two diagonal subunits are shown for clarity. Residues at the pore constrictions are shown in sticks. f, Pore profile of hTRPC3 in the open state (T561A-C4) calculated using HOLE2. g, Pore profile of srTRPM2 in the open state (PDB ID: 8SRD) calculated using HOLE2.

**Figure 3.**
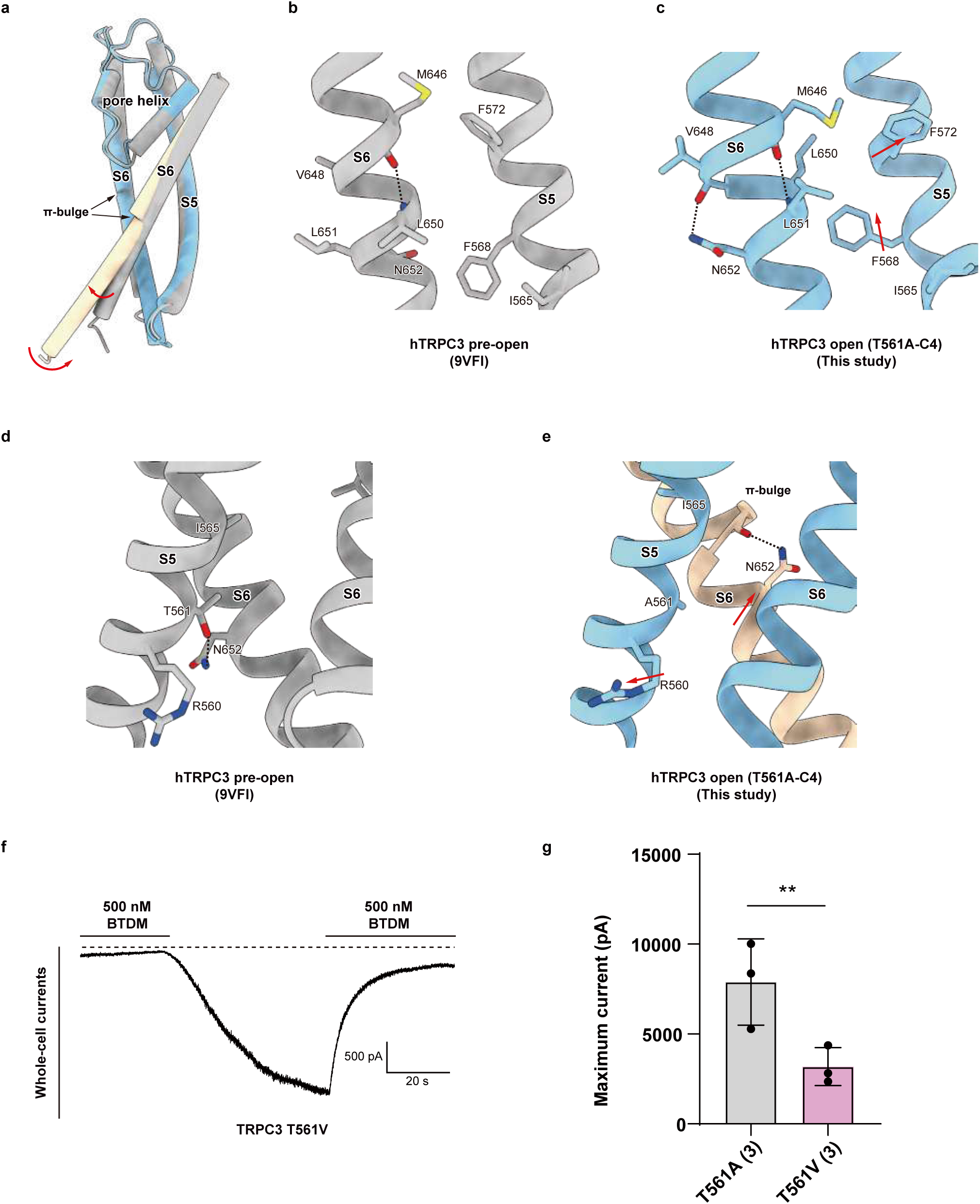
Conformational changes of hTRPC3 from pre-open to open state. a, Structural comparison of hTRPC3 between the open state (T561A-C4, colored) and the pre-open state (gray, PDB ID: 9VFI). The rotation and tilting movement of the lower half of S6 are denoted with arrows. The π-bulge of the hTRPC3 in the open state near the middle of S6 is marked. b, The close-up view of hTRPC3 in the pre-open state. The hydrogen bond is denoted with dashes. c, The close-up view of hTRPC3 in the open state. The hydrogen bond is denoted with dashes. d, The close-up view of hTRPC3 in the pre-open state. The hydrogen bond is denoted with dashes. e, The close-up view of hTRPC3 in the open state. The hydrogen bond is denoted with dashes. f, Whole-cell currents of the hTRPC3 T561V mutant recorded at −60 mV. The dashed line denotes the position of zero current. The recording has been repeated three times with similar results. g, Statistics of the constitutive whole cell currents recorded at −60 mV. The data are shown as the mean ± SD. The number of replicates in each group is indicated in parentheses. The data were analyzed using t-tests with p-values indicated above the corresponding groups.

The image processing workflow yielded two major structures, which share an identical CTD (Supplementary Fig. 3k) but differ in their TMD. One structure exhibits canonical C4 symmetry, resolved at 2.93 Å resolution (referred to as T561A-C4, Supplementary Fig. 2-3), while the other exhibits C2 symmetry, resolved at 3.16 Å resolution (referred to as T561A-C2, Supplementary Fig. 2-3). We will first discuss the C4 structure (Fig. 2-4), which represents the open state, and then address the C2 structure (Fig. 5), which is a non-conductive state.

**Figure 4.**
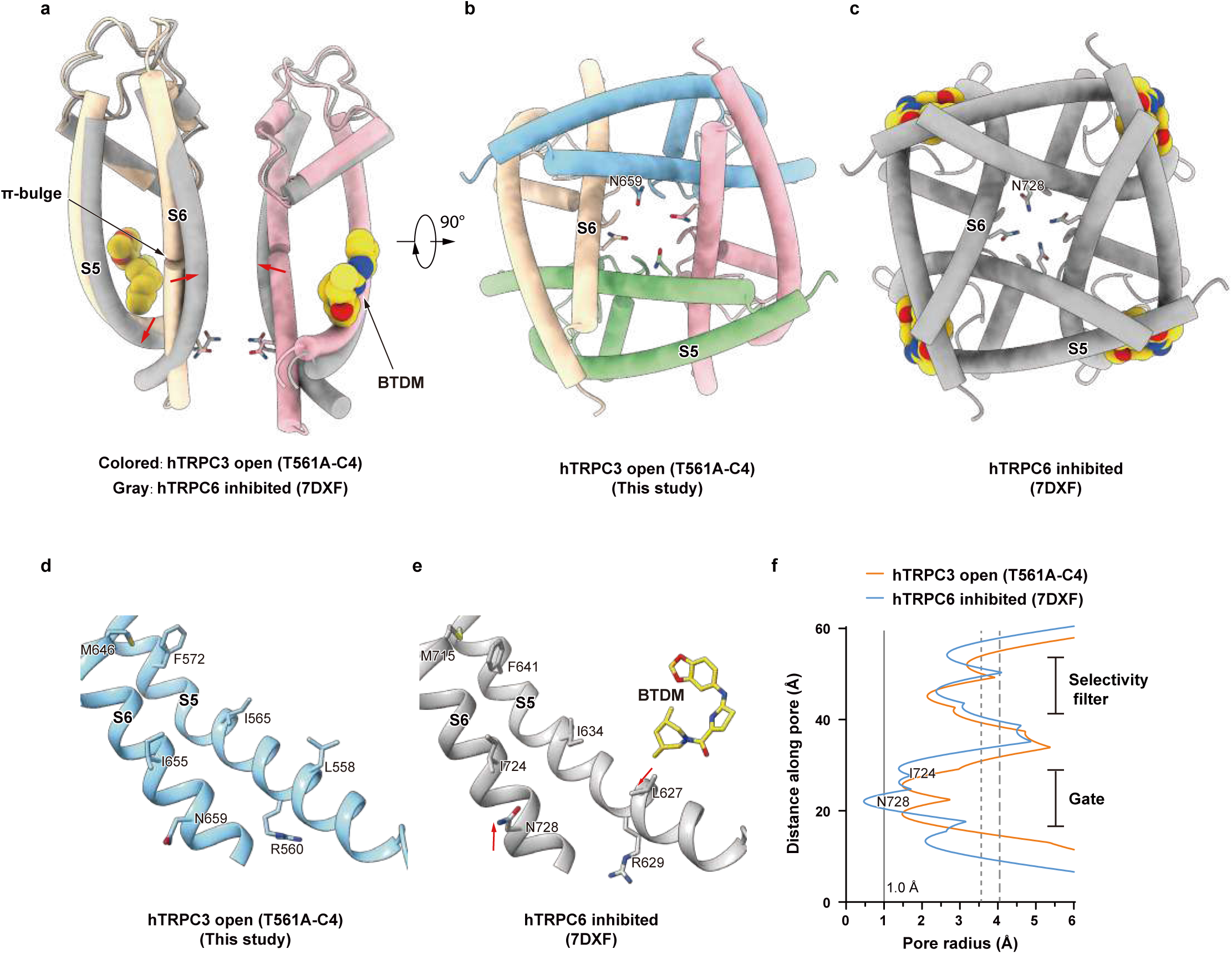
Conformational changes of hTRPC3 from open to BTDM-inhibited state. a, Structural comparison between hTRPC3 in the open state (T561A-C4, colored) and hTRPC6 in the BTDM-inhibited closed state (gray, PDB ID: 7DXF). The movement of S6 is denoted with arrows. The π-bulge near the middle of S6 is marked. BTDM is shown as spheres. b, a 90°-rotated bottom view of a, showing the open gate of hTRPC3 in the open state. c, a 90°-rotated bottom view of a, showing the closed gate of hTRPC6 in the BTDM-inhibited state. d, The close-up view of hTRPC3 in the open state. e, The close-up view of hTRPC6 in the BTDM-inhibited closed state. f, Pore radius calculated using HOLE2. The gray solid line denotes the radius of the dehydrated calcium ion (1.0 Å). The gray dashed lines denote the radius of the hydrated Na^+^ (dotted) and Ca^2+^ ions (dashed).

**Figure 5.**
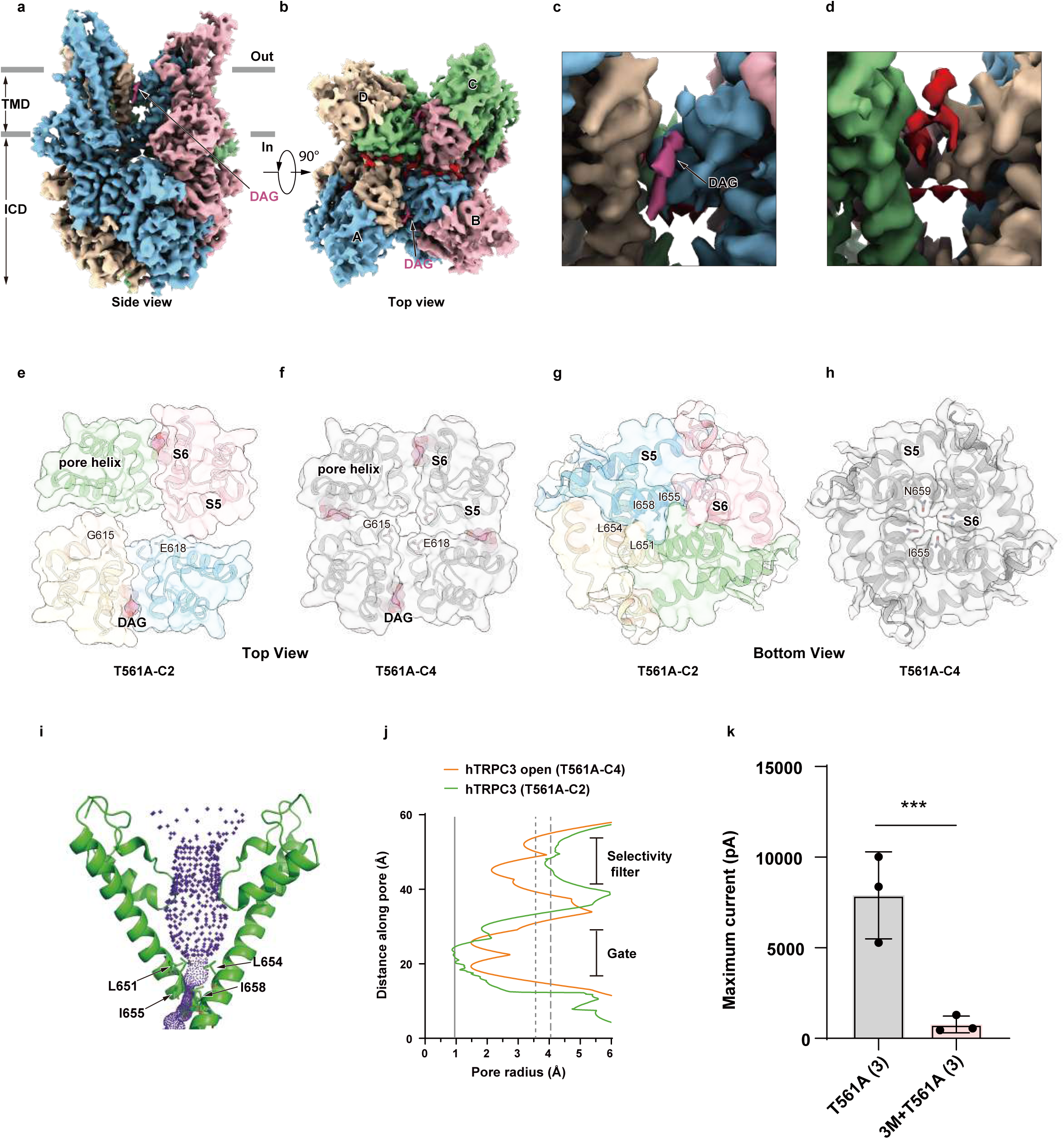
DAG binding stabilizes the open state. a, Cryo-EM density map of the hTRPC3 T561A mutant (T561A-C2) shown in side view. Subunits A, B, C, and D are colored the same as Fig. 1d. DAG is colored in purple. b, a 90°-rotated top view of a. c, Close-up view of panel b at the intact L2 DAG binding site between subunit A-D and B-C. d, Close-up view of panel b at the disrupted L2 DAG binding site between subunit A-B and C-D. e, Top view of the extracellular surface of the T561A-C2 structure. DAG is shown as spheres. f, Top view of the extracellular surface of the T561A-C4 structure. DAG is shown as spheres. g, Bottom view of the closed gate of T561A-C2 structure. h, Bottom view of the open gate of T561A-C4 structure. i, Pore profile of hTRPC3 T561-C2 structure calculated using HOLE2. Only the S6 helices of two diagonal subunits are shown for clarity. Residues at the pore constrictions are shown in sticks. j, Pore radius calculated using HOLE2. The gray solid line denotes the radius of the dehydrated calcium ion (1.0 Å). The gray dashed lines denote the radius of the hydrated Na^+^ (dotted) and Ca^2+^ ions (dashed). k, Statistics of the constitutive whole cell currents recorded at −60 mV. The data are shown as the mean ± SD. The number of replicates in each group is indicated in parentheses. The data were analyzed using t-tests with p-values indicated above the corresponding groups. The data of T561A is the same as shown in Fig. 3g.

The T561A-C4 structure shows a pronounced dilation of the pore compared to the pre-open state (Fig. 2d-f). Pore profile calculations using HOLE2 show that the radius at the narrowest constriction increases from 0.8 Å to 1.5 Å (Fig. 2d), which might be sufficient to allow permeation of partially hydrated calcium ions. In this conformation, the pore is lined by the side chains of the hydrophobic residue I655 and the hydrophilic residue N659 of hTRPC3 (Fig. 2f), which is different from those in the pre-open state (Fig. 2e). The pore radius and pore-lining residues are similar to those observed in the open state of TRPM2 (PDB ID: 8SRD, Fig. 2g) ^17^. Therefore, we refer to it as the agonist-bound open state structure of hTRPC3.

### Structural transitions during hTRPC3 opening

In the T561A-C4 structure, the L2 site is occupied by DAG (Fig. 2c). A comparison of this open-state structure with the DAG-bound pre-open state (PDB ID: 9VFI) ^13^ reveals that their CTDs are identical (Supplementary Fig. 3l); thus, we focused our analysis on the TMD, in which the major structural differences are observed in the S5 and S6 helices (Supplementary Video 1).

In the pre-open state, the S6 helix adopts a canonical α-helical conformation (Fig. 3a-b). The carbonyl group of M646 made a hydrogen bond with the amino group of L650 (Fig. 3b). N652 on S6 forms a polar interaction with T561 on S5 (Fig. 3d). However, in the T561A mutant, this N652-T561 interaction is disrupted (Fig. 3e). To test whether the T561A mutation activates the channel by disrupting this specific interaction, we introduced a T561V mutation into the wild-type hTRPC3. Valine is similar in size to threonine but cannot form a polar interaction with N652. The T561V mutant also exhibited robust constitutive currents (Fig. 3f-g), supporting the conclusion that the activation mechanism mainly involves the disruption of the N652-T561 interaction.

In the T561A-C4 structure, a π-bulge structure is formed, and the carbonyl group of M646 made a hydrogen bond with the amino group of L651 (Fig. 3c). N652 instead forms a new polar interaction with the main-chain carbonyl group of V648, stabilizing the π-bulge (Fig. 3c). Consequently, the lower half of S6 undergoes not only a rotation but also a tilting movement (Fig. 3a). The rotation of S6 reconfigures the pore lining residues (Fig. 2e-f) and local interactions with the adjacent S5 helix (Fig. 3d-e). This leads to concomitant conformational changes in S5, including the rotation of side chains of residues R560, I565, F568, and F572 (Fig. 3b-e). The tilting of the S6 helix results in pore dilation and thus channel opening.

### Mechanism of BTDM inhibition

BTDM is a high-affinity inhibitor of both hTRPC3 and hTRPC6 ^9^. We have previously determined the structure of hTRPC6 in complex with BTDM at 2.9 Å resolution and identified its detailed binding mode ^11^. Given the high sequence similarity between hTRPC3 and hTRPC6, it is likely that BTDM binds and inhibits hTRPC3 in a similar manner. To understand how BTDM inhibits the hTRPC3 T561A mutant, we compared the T561A-C4 structure with the hTRPC6-BTDM complex structure (Fig. 4 and Supplementary Video 2). This comparison suggests that upon BTDM binding, the 3,5-dimethylpiperidin group of BTDM would effectively push L558 on S5 of hTRPC3 toward the pore center (Fig. 4d-e). Because of the intimate interactions between S5 and the adjacent gating helix S6, S6 consequently moves inward, leading to pore closure by the constrictions at N659 (Fig. 4d-f). Notably, in the BTDM-inhibited closed state, the π-bulge and pore-lining residues are preserved compared with the open state ^11^.

### DAG stabilizes the open state

In the T561A-C2 structure (Fig. 5a-b), we observed that two diagonal L2 sites (at the A-D and B-C subunit interfaces) are occupied by DAG (Fig. 5c), while the other two L2 sites (at the A-B and C-D interfaces) are disrupted and free of DAG density (Fig. 5d). As a result, the subunit interfaces associated with the unoccupied sites are misaligned (Fig. 5e-f). This misalignment leads to a distortion of the selectivity filter (Fig. 5e-f) and a rectangular gate at the bundle crossing where L651, L654, I655, and I658 blocks the pore (Fig. 5g-h). Pore profile calculations confirm that this configuration is non-conductive (Fig. 5i-j). These observations suggest that the DAG binding at the L2 site stabilizes the subunit interface, thereby maintaining the open state of the hTRPC3 channel. Consistent with this model, we found that decreasing the DAG binding by combining the 3M mutations (E603A, V637A, and K607A together) on the hTRPC3 T561A mutant greatly decreased its constitutive currents (Fig. 5k).

## Discussion

By comparing the agonist-free resting state (TRPC3-3M), the DAG-bound pre-open state, and the DAG-bound open state (T561A-C4), we did not observe large-scale conformational changes in the binding site of the glycerol head group of DAG (Supplementary Fig. 4a-c). However, the binding site of the two acyl tails of DAG, is in close proximity to residues that undergo substantial rearrangements during channel opening, including F572, M569, and I565 on S5, and V649 and L650 on S6 (Supplementary Fig. 4d). Moreover, DAG binding appears to stabilize the open state, as the T561A-C2 structure shows that the loss of DAG at two subunit interfaces results in a distorted, non-conductive pore (Fig. 5). Furthermore, abolishing the DAG-binding site by 3M mutations dramatically decreased the currents (Fig. 5k). Together, these observations support the role of DAG as an agonist that stabilizes the open conformation of hTRPC3.

We also observed differential effects of hTRPC3 inhibitors on the T561A mutant. BTDM binds at the Inhibitor Binding Pocket-C (IBP-C) site ^9,11^, which is located between the pore and voltage sensor-like domain (VSLD). BTDM effectively inhibits the constitutive current of the T561A mutant (Fig. 2a). In contrast, SAR7334, which binds at the Inhibitor Binding Pocket-A (IBP-A) site in the VSLD ^11^, does not inhibit the currents of the T561A mutant (Fig. 2a). This correlates with the fact that the VSLD of hTRPC3 does not show conformational differences between the pre-open and the open state structures (Supplementary Fig. 4e) and suggests that the VSLD regulates hTRPC3 activity at the upstream of the T561 site.

Leveraging the structural information presented here, along with the BTDM-bound closed-state structure of the closely related hTRPC6, we propose a hypothetical gating trajectory for hTRPC3. This trajectory encompasses transitions from the resting state to the pre-open state, followed by agonist-stabilized opening, and finally to a BTDM-inhibited state (Supplementary Video 3). Notably, we observed an α-to-π transition in the middle of the S6 helix, which leads to rotation and tilting of S6 and subsequent pore opening of hTRPC3. This correlates with a global structural survey of TRP channel gating mechanisms, in which researchers found that the α-to-π transition of the S6 helix has also been associated with activation in other TRP channel families ^18^. Given the high sequence homology and likely conserved gating mechanism among TRPC3, TRPC6, and TRPC7, we speculate that this gating trajectory may represent a common mechanism for this subfamily.

## Methods

### Cell Culture

Sf9 insect cells (Thermo Fisher Scientific) were cultured in SIM SF (Sino Biological) at 27 °C. Adapted Expi293F cells (donated by Shanghai OPM Biosciences) were cultured in 293F Hi-exp Medium (Shanghai OPM Biosciences) at 37 °C with 6% CO_2_ and 70% humidity. FreeStyle 293F suspension cells (Thermo Fisher Scientific) were cultured in FreeStyle 293 medium (Thermo Fisher Scientific) supplemented with 1% fetal bovine serum (FBS, VisTech), 67 μg/mL penicillin (Macklin), and 139 μg/mL streptomycin (Macklin) at 37 °C with 6% CO_2_ and 70% humidity. The cell lines were routinely checked to be negative for mycoplasma contamination, but have not been authenticated.

### Constructs

The hTRPC3 constructs were cloned into a modified N-terminal GFP-tagged BacMam vector for electrophysiology assay as described previously ^9^. For protein purification, the hTRPC3 constructs were inserted into an N-terminal GFP-MBP (Maltose Binding Protein)-tagged BacMam vector.

### Electrophysiology

For whole-cell recordings of TRPC3, the pipette solution was 120 CsMe (Cesium methanesulfonate), 25 CsCl, 2 MgCl_2_, 2 EGTA, and 30 HEPES (pH = 7.4 adjusted by CsOH) in mM. The bath solution contains 140 NaCl, 5 CsCl, 2 MgCl_2_, 1 EDTA, 2 EGTA, 10 Glucose, and 10 HEPES (pH = 7.4, adjusted by NaOH) in mM.

The concentrations of 1-Oleoyl-2-acetyl-sn-glycerol (OAG, Sigma, O6754), BTDM (donated by Dizal Pharmaceutical) used in patch-clamp are marked out in the figures.

Patch electrodes were pulled by a horizontal microelectrode puller (P-1000, Sutter Instrument Co., USA) to a tip resistance of 2.0–4.0 MΩ. An MPS-2 perfusion system (Yibo Company, Wuhan, China) was used for buffer change. The TRPC currents were recorded at a holding potential of −60 mV through an Axopatch 200B amplifier (Axon Instruments, USA). The currents with amplitudes greater than 150 pA were used for further analysis. Data were further low-pass filtered by pCLAMP 10.4 software. The current traces were plotted using Origin 8. Peak currents were statistically analyzed in GraphPad Prism 8.

### Protein expression and purification

For expression of hTRPC3 T561A mutant, adapted Expi293F cells were cultured in 293F Hi-exp Medium (Shanghai OPM Biosciences) at a density of 3.0 × 10^6^ mL^−1^ and were transfected with the hTRPC3 T561A mutant plasmid using polyethyleneimine (PEI MAX, Polysciences). 10 mM sodium butyrate and 400 nM BTDM were added to the medium 12 h post-transfection, and the cells were cultured for another 48 h before harvesting. For harvesting, 4 mM EDTA was added to block divalent cations in the culture medium, and cells were washed four times using 1 mM EDTA in TBS buffer (20 mM Tris pH 8.0 at 4 °C, 150 mM NaCl) at room temperature to remove most of the BTDM before being frozen at −80 °C.

For purification of the hTRPC3 T561A mutant, 293F cells were grown in FreeStyle 293 medium. Cell pellets corresponding to 800 mL culture were solubilized in 50 mL lysis buffer [2 mg/mL 4F (DWFKAFYDKVAEKFKEAF), 25 mM Tris pH 8.0 at 4 °C, 5 mM DTT, 1 mM EDTA, and protease inhibitors] and rotated at 4 °C for 1 h. After centrifugation at 40,000 rpm for 40 min in the Type 50.2 rotor (Beckman), the supernatant was loaded onto a column packed with amylose resin (NEB). The resin was washed with TBS buffer containing 5 mM DTT and 1 mM EDTA. Protein was eluted with MBP elution buffer (80 mM maltose, 100 mM NaCl, 20 mM Tris pH 8.0 at 4 °C, 5 mM DTT, and 1 mM EDTA) and incubated with H3CV protease at 4 °C overnight. The next day, the protein was incubated with MSP2N2 at a 1:10 M ratio for 1 h at room temperature, then the mixture was loaded onto Streptactin resin and washed with TBS containing 5 mM DTT. The target protein was eluted with Strep elution buffer (50 mM Tris pH 8.0 at 4 °C, 100 mM NaCl, 5 mM desthiobiotin, and 5 mM DTT) and loaded onto Superose-6 increase (GE Healthcare) running with S6 buffer (20 mM HEPES pH 7.5, 100 mM NaCl, and 1 mM TCEP) for further purification. The peak fractions that contain the tetrameric hTRPC3 channel in nanodiscs were combined and concentrated for cryo-EM studies.

For expression of hTRPC3 3M mutant, HEK293F cells grown in FreeStyle 293 medium at 37 °C with a density of 2.3 × 10^6^ mL^-1^ were infected by BacMam viruses. 10 mM sodium butyrate was added 8–12 h post-infection, and the temperature was lowered to 30 °C. Cells were collected 48 h post-infection and washed twice with TBS buffer before being frozen at −80 °C.

For purification of hTRPC3 3M mutant, cell pellets corresponding to 800 mL culture were solubilized in 50 mL lysis buffer (2 mg/mL 4F, 25 mM Tris pH 8.0 at 4 °C, 5 mM DTT, 1 mM EDTA, and protease inhibitors) and rotated at 4 °C for 1 h. After centrifugation at 40,000 rpm for 40 min in the Type 50.2 rotor (Beckman), the supernatant was loaded onto a column packed with amylose resin (NEB). The resin was washed with TBS buffer containing 5 mM DTT and 1 mM EDTA. Protein was eluted with MBP elution buffer (80 mM maltose, 100 mM NaCl, 20 mM Tris pH 8.0 at 4 °C, 5 mM DTT, and 1 mM EDTA) and incubated with MSP2N2 at a 1:10 M ratio at 4 °C overnight. And then the mixture was loaded onto Streptactin resin and washed with TBS containing 5 mM DTT. The target protein was eluted with Strep elution buffer (50 mM Tris pH 8.0 at 4 °C, 100 mM NaCl, 5 mM desthiobiotin, and 5 mM DTT) and incubated with H3CV protease at 4 °C overnight. The next day, the protein was loaded onto a GST column, and the wash and FT fractions were collected using TBS (pH 7.5 at RT) supplemented with 5 mM DTT to remove H3CV protease. Then the mixture was loaded onto Superose-6 increase (GE Healthcare) running with S6 buffer (20 mM HEPES pH 7.5, 100 mM NaCl, and 1 mM TCEP) for further purification. The peak fractions that contain the tetrameric hTRPC3 channel in nanodiscs were combined and concentrated for cryo-EM studies.

### Cryo-EM sample preparation

1 mM CaCl_2_ was added to the purified hTRPC3 T561A mutant with A280 ≈ 0.12. After centrifuging at 40,000 rpm for 30 min, the supernatant was collected and supplemented with 0.5 mM fluorinated octyl maltoside (FOM). The protein was added on Quantifoil (1.2/1.3) gold grids coated with graphene ^19^ for cryo-EM sample preparation.

1 mM CaCl_2_ was added to the purified hTRPC3 3M mutant with A280 ≈ 1.6. After centrifuging at 40,000 rpm for 30 min, the supernatant was collected and supplemented with 0.5 mM fluorinated octyl maltoside (FOM). The protein was added onto GIG (1.2/1.3) Cu grids coated with graphene ^19^ for cryo-EM sample preparation.

### Cryo-EM data collection

Cryo-EM grids were screened on the Talos Arctica electron microscope (Thermo Fisher Scientific) operating at 200 kV using a K2 camera (Thermo Fisher Scientific). The screened grids of hTRPC3 T561A mutant were then transferred to the 300 kV Titan Krios (Thermo Fisher Scientific) with a K3 camera (Gatan) and an energy filter set to a slit width of 20 eV at a magnification of 47,000×with a pixel size of 1.067 Å, and the defocus ranging from −1.5 to −1.8 μm, with a total dose about 50 e^−^/Å^2^ and a dose rate of 22.36 e^−^/pixel/s on detector.

The screened grids of hTRPC3 3M mutant were then transferred to the 300 kV Titan Krios (Thermo Fisher Scientific) with a K3 camera (Gatan) and an energy filter set to a slit width of 20 eV at a magnification of 60,000× with a pixel size of 0.833 Å, and the defocus ranging from −1.5 to −1.8 μm, with a total dose about 50 e^−^/Å^2^ and a dose rate of 22.15 e^−^/pixel/s on detector.

### Cryo-EM data processing

The hTRPC3 T561A mutant datasets were first gain-corrected, motion-corrected, anisotropic magnification corrected, and dose-weighted by MotionCor2-1.3.2 ^20^. The selected micrographs were then imported into cryoSPARC-4.5.3 ^21^. A round of template picking, 2D and 3D classification, and 3D refinement was performed in cryoSPARC-4.5.3. The resulting particles from the best class were used as “Seeds” for seed-facilitated 3D classification ^22^ of particles extracted by Topaz ^23^. The particles selected after multi-reference 3D classification were divided into two groups, which were subsequently used for final non-uniform refinement ^24^ with C4 and C2 symmetry, respectively. The resolution estimation was based on the gold standard FSC 0.143 cut-off.

The data processing procedures for the hTRPC3 3M mutant followed the same strategy as described above. The selected micrographs were then imported into cryoSPARC-4.5.3. A round of template picking, 2D and 3D classification, and 3D refinement was performed in cryoSPARC-4.5.3. The resulting particles from the best class were used as “Seeds” for seed-facilitated 3D classification of particles extracted by Topaz. The particles selected after multi-reference 3D classification were subsequently used for final non-uniform refinement with C4 symmetry. The resolution estimation was based on the gold standard FSC 0.143 cut-off.

### Model building, refinement, and validation

The hTRPC3 models were manually rebuilt using Coot-0.9.8.93 ^25^. based on the published models 7DXC and further refined using PHENIX1.19.2 ^26,27^.

### Quantification and statistical analysis

Data processing and statistical analysis were conducted using the software GraphPad Prism. Statistical details could be found in the methods details and figure legends. Global resolution estimations of cryo-EM density maps are based on the 0.143 Fourier Shell Correlation criterion ^28^. The local resolution was estimated using cryoSPARC. The number of independent experiments (N) and the relevant statistical parameters for each experiment (such as mean or standard deviation) are described in the figure legends. No statistical methods were used to pre-determine sample sizes.

## Data Availability

The data that support the findings of this study are available from the corresponding author upon request. Cryo-EM maps and atomic coordinates of hTRPC3 3M mutant in the resting state (hTRPC3-3M) have been deposited in the EMDB and PDB database under the ID codes EMDB: EMD-80010 and PDB: 25CG, respectively. Cryo-EM maps and atomic coordinates of hTRPC3 T561A mutant in the open state (hTRPC3-T561A-C4) have been deposited in the EMDB and PDB database under the ID codes EMDB: EMD-80011 and PDB: 25CH, respectively. Cryo-EM maps and atomic coordinates of hTRPC3 T561A mutant in the distorted closed state with C2 symmetry (hTRPC3-T561A-C2) have been deposited in the EMDB and PDB database under the ID codes EMDB: EMD-80012 and PDB: 25CI, respectively.

## Acknowledgments

We thank all of the Chen Lab members for their kind help. We thank Dr. Xiaolin Zhang at Dizal Pharmaceutical Company for providing BTDM compound. Cryo-EM data collection was supported by the Electron microscopy laboratory, and the Cryo-EM platform of Peking University. Part of the structural computation was also performed on the Computing Platform of the Center for Life Science and High-performance Computing Platform of Peking University. We thank the National Center for Protein Sciences at Peking University in Beijing, China, for assistance with negative stain EM. The work is supported by grants from the Ministry of Science and Technology of China (National Key R&D Program of China, 2022YFA0806504 to L.C.), National Natural Science Foundation of China (32225027 to L.C., 52021006 to H.P., 32301009 to W.G.), and the Center for Life Sciences (CLS to L.C.).

## Author contributions

L.C. initiated the project. J.Z., and Y.T. purified the protein, prepared the cryo-EM sample, collected the cryo-EM data, and built the model. Y.T., Y.C., and W.G. carried out electrophysiological experiments. P.H. and Z.X. prepared the graphene-coated grids. All authors contributed to the manuscript preparation.

---

**Supplementary Video 1 | Conformational changes of hTRPC3 from the pre-open to the open state.** hTRPC3 is colored the same as shown in Fig. 1d. Morphs of hTRPC3 between the pre-open state (PDB ID: 9VFI) and the open state (T561A-C4) are generated using Chimera X.

**Supplementary Video 2 | Conformational changes of hTRPC3 from the open state to the BTDM-inhibited state.** hTRPC3 is colored the same as shown in Fig. 1d. The structure of hTRPC3 in the BTDM inhibited state was generated with SWISS-MODEL using the structure of hTRPC6 in complex with BTDM (PDB ID: 7DXF) as the template. BTDM is shown as spheres. Morphs of hTRPC3 between the open state (T561A-C4) and the BTDM-inhibited state are generated using Chimera X.

**Supplementary Video 3 | A hypothetical gating trajectory of hTRPC3.** The morphs of hTRPC3 from the resting state (hTRPC3-3M) to the pre-open state (PDB ID: 9VFI), followed by agonist-stabilized open state (T561A-C4), and finally to a BTDM-inhibited state (modelled from PDB ID: 7DXF).

**Supplementary Figure 1.**
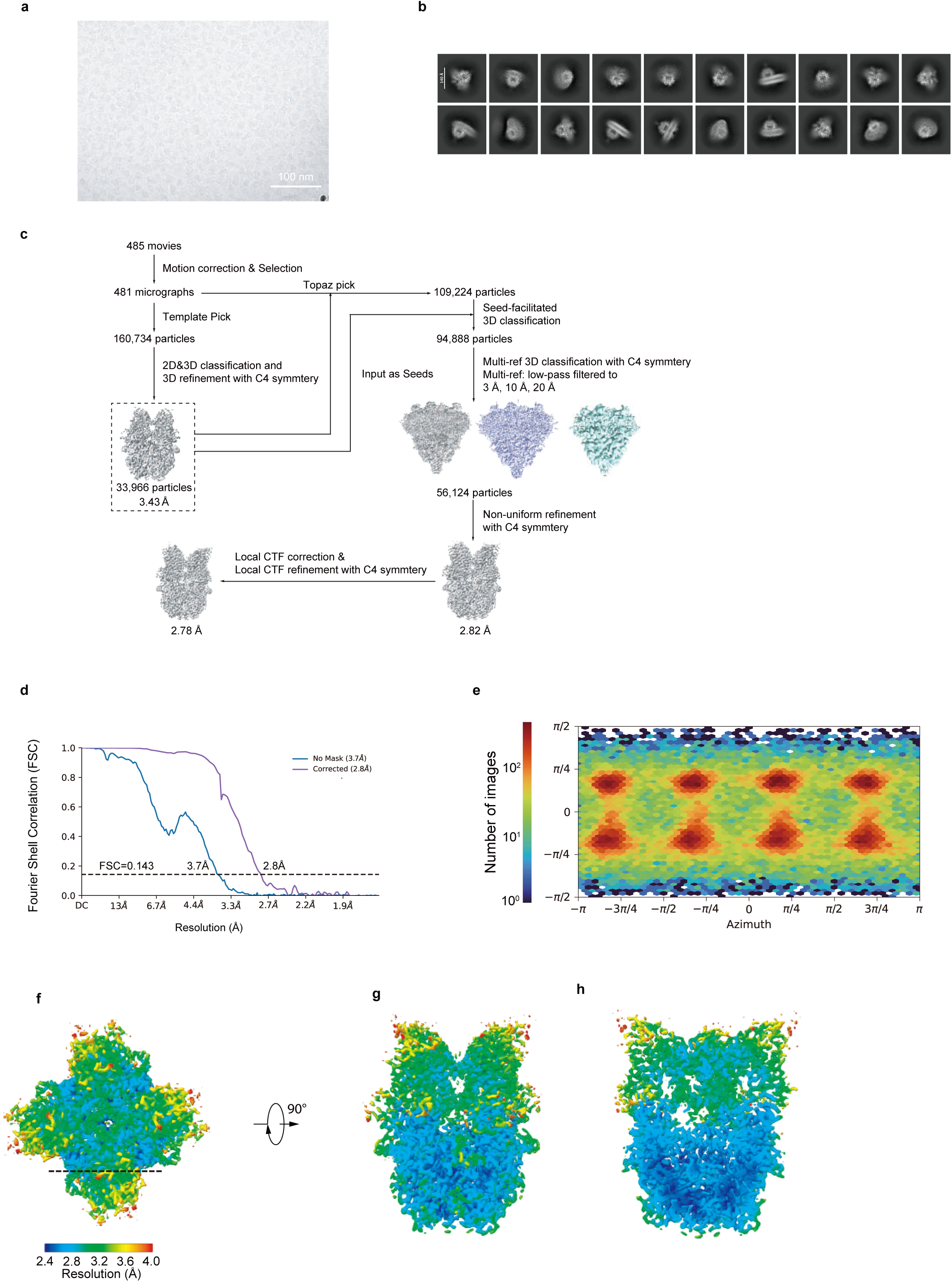
Cryo-EM image analysis of hTRPC3 3M. a, Representative raw micrograph of hTRPC3 3M purified using 4F-disc. b, Representative 2D class average of hTRPC3 3M. c, Flowchart of hTRPC3 3M cryo-EM data processing. d, FSC curves of final refinement for hTRPC3 3M after correction for masking effects. Resolution estimations were based on the criterion of the FSC 0.143 cutoff. e, Angular distribution of final refinement for hTRPC3 3M. This is a standard output from cryoSPARC. f, Local resolution estimation of hTRPC3 3M in top view. g, Local resolution estimation of hTRPC3 3M in side view. h, The cross-section of local resolution estimation of hTRPC3 3M. The position of the cross-section was indicated as the dashed line in f.

**Supplementary Figure 2.**
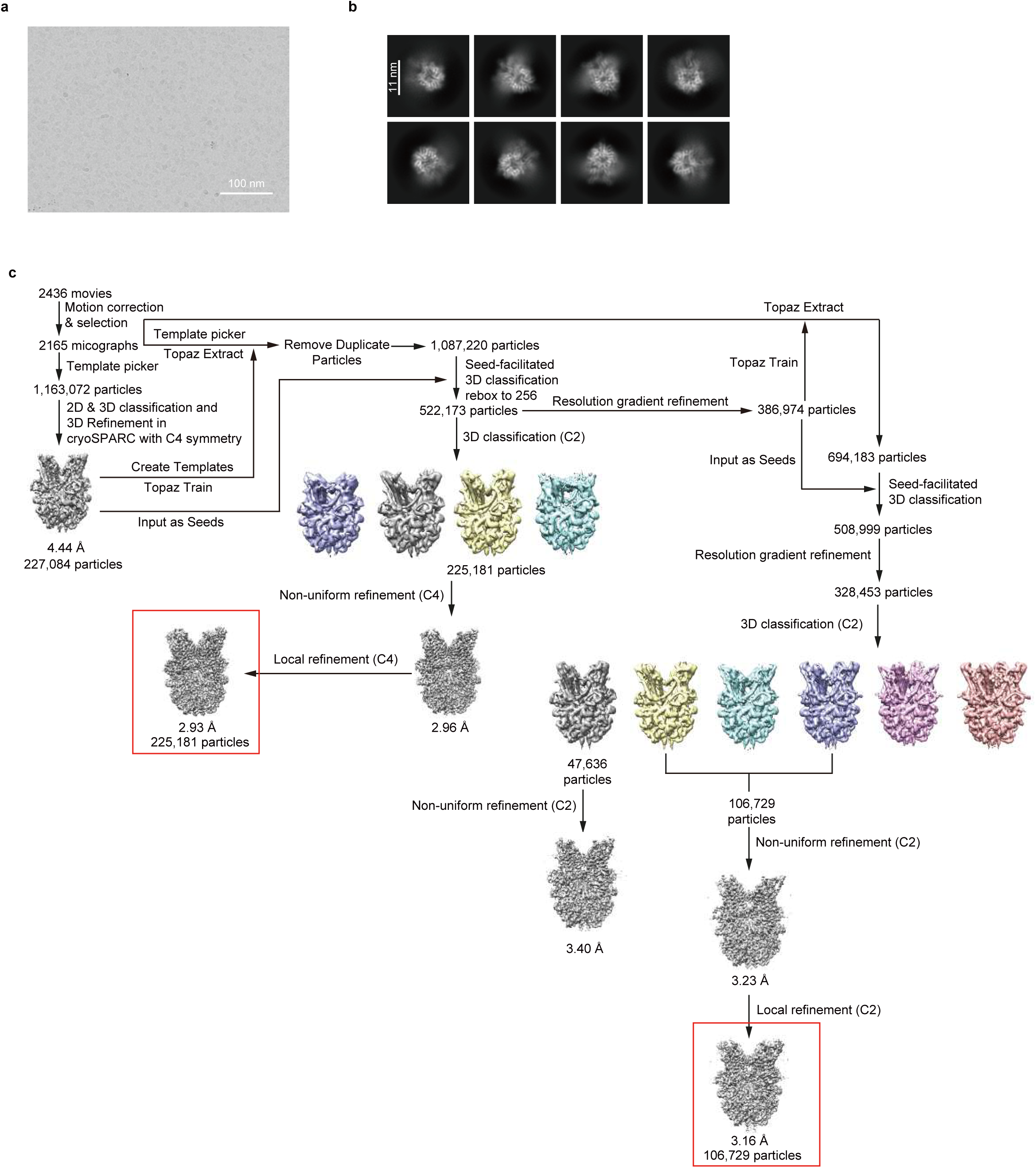
Cryo-EM image analysis of hTRPC3 T561A mutant. a, Representative raw micrograph of hTRPC3 T561A purified using 4F-disc. b, Representative 2D class average of hTRPC3 T561A. c, Flowchart of hTRPC3 T561A cryo-EM data processing.

**Supplementary Figure 3.**
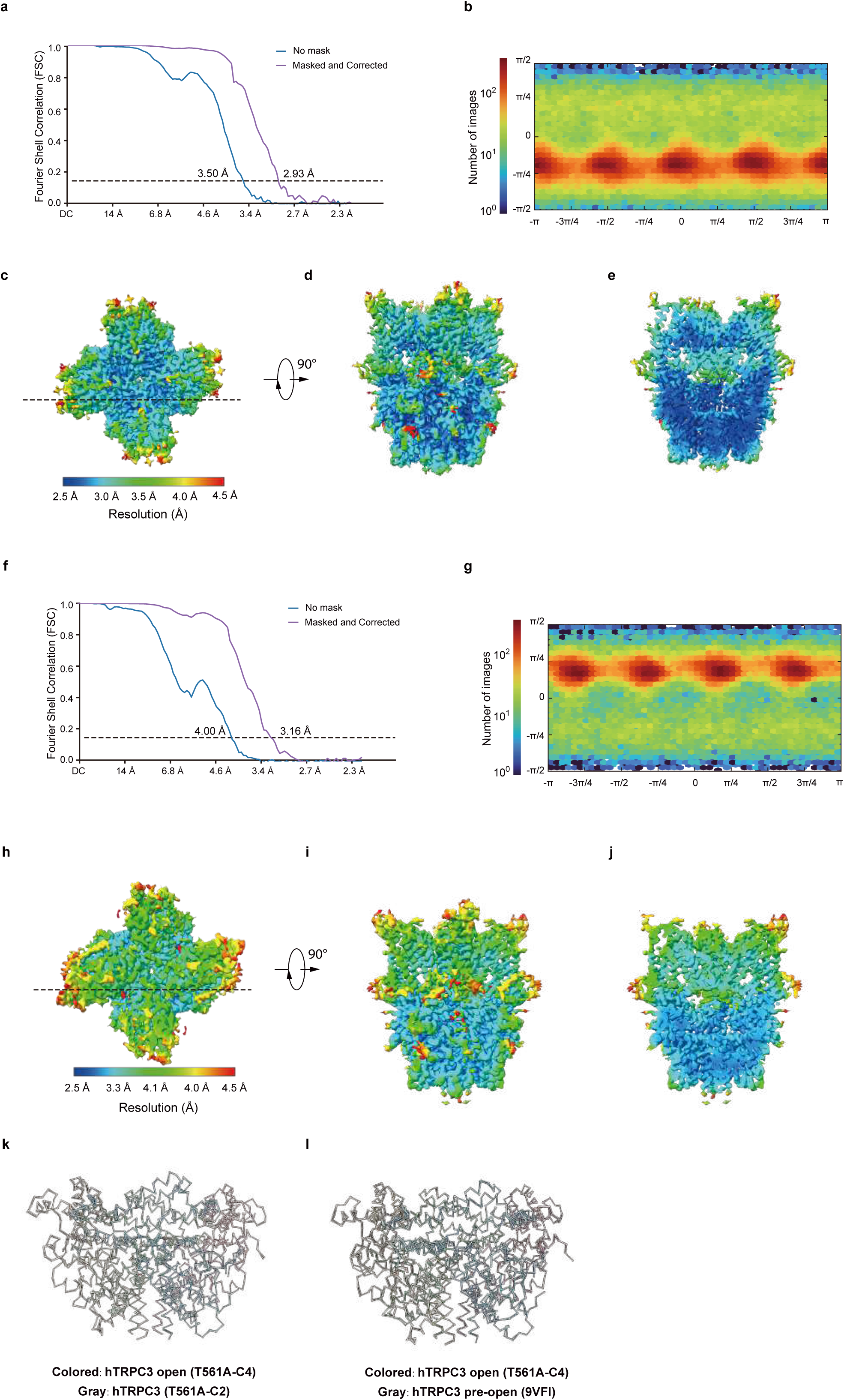
Cryo-EM reconstruction of hTRPC3 T561A mutant. a, FSC curves of final refinement for hTRPC3 T561A-C4 after correction for masking effects. Resolution estimations were based on the criterion of the FSC 0.143 cutoff. b, Angular distribution of final refinement for hTRPC3 T561A-C4. This is a standard output from cryoSPARC. c, Local resolution estimation of hTRPC3 T561A-C4 in top view. d, Local resolution estimation of hTRPC3 T561A-C4 in side view. e, The cross-section of local resolution estimation of hTRPC3 T561A-C4. The position of the cross-section was indicated as the dashed line in c. f, FSC curves of final refinement for hTRPC3 T561A-C2 after correction for masking effects. Resolution estimations were based on the criterion of the FSC 0.143 cutoff. g, Angular distribution of final refinement for hTRPC3 T561A-C2. This is a standard output from cryoSPARC. h, Local resolution estimation of hTRPC3 T561A-C2 in top view. i, Local resolution estimation of hTRPC3 T561A-C2 in side view. j, The cross-section of local resolution estimation of hTRPC3 T561A-C2. The position of the cross-section was indicated as the dashed line in h. k, The structural comparison of CTD between hTRPC3 T561A-C4 and hTRPC3 T561A-C2 (gray). l, The structural comparison of CTD between hTRPC3 T561A-C4 and wild type hTRPC3 (gray, PDB ID: 9VFI).

**Supplementary Figure 4.**
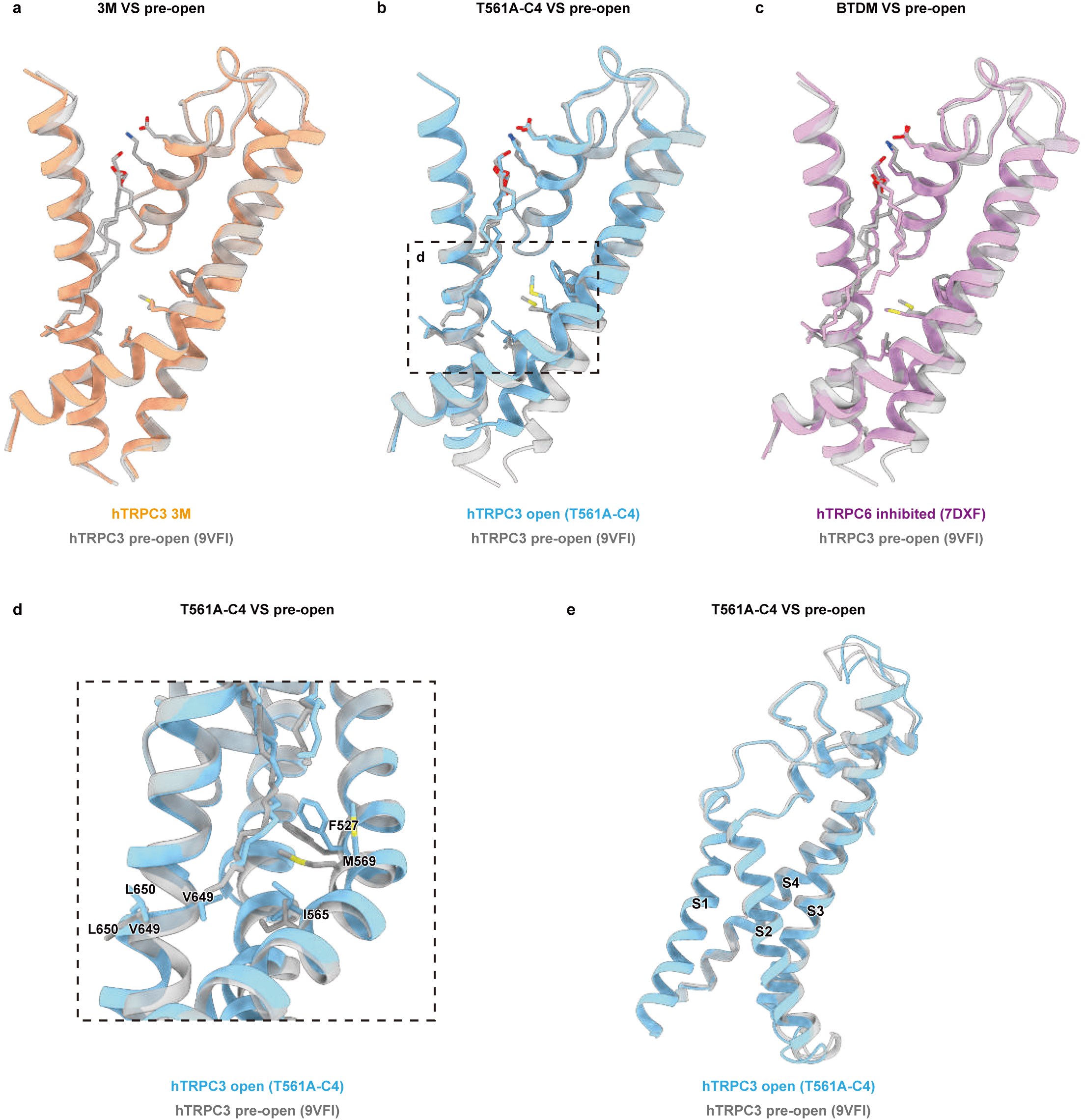
Structural comparisons. a, Structural comparison between apo-resting hTRPC3 (hTRPC3-3M, orange) and DAG-bound pre-open hTRPC3 (wild-type hTRPC3, PDB ID: 9VFI gray) at the DAG-binding site. DAG is shown as sticks. b, Structural comparison between open hTRPC3 (hTRPC3 T561A-C4, blue) and DAG-bound pre-open hTRPC3 (wild-type hTRPC3, PDB ID: 9VFI, gray) at the DAG-binding site. c, Structural comparison between BTDM-inhibited hTRPC6 (PDB ID: 7DXF, purple) and DAG-bound pre-open hTRPC3 (wild-type hTRPC3, PDB ID: 9VFI, gray) at the DAG-binding site. d, Close-up view of the DAG binding site of panel b near its tail. Conformational changes of side chains during channel opening are denoted with arrows. e, Structural comparison open hTRPC3 (hTRPC3 T561A-C4, blue) and DAG-bound pre-open hTRPC3 (wild-type hTRPC3, PDB ID: 9VFI, gray) at VSLD.

**Supplementary Table S1.** Cryo-EM data collection, refinement and validation statistics.

| PDB ID<br>EMDB ID | hTRPC3-3M<br>25CG<br>EMD-80010 | T561A-C4<br>25CH<br>EMD-80011 | T561A-C2<br>25CI<br>EMD-80012 |
| --- | --- | --- | --- |
| **Data collection and processing** |  |  |  |
| Magnification | 60,000 × | 47,000 × | 47,000 × |
| Voltage (kV) | 300 | 300 | 300 |
| Electron exposure (e<sup>-</sup>/Å<sup>2</sup>) | 50 | 50 | 50 |
| Defocus range (μm) | -1.5 to -1.8 | -1.5 to -1.8 | -1.5 to -1.8 |
| Pixel size (Å) | 0.833 | 1.067 | 1.067 |
| Symmetry imposed | *C4* | *C4* | *C2* |
| Initial particle images (no.) | 160,734 | 1,163,072 | 1,163,072 |
| Final particle images (no.) | 56,124 | 225,181 | 106,729 |
| Map resolution (Å) | 2.78 | 2.93 | 3.16 |
| FSC threshold | 0.143 | 0.143 | 0.143 |
| Map resolution range (Å) | 250-2.78 | 250-2.93 | 250-3.16 |
| **Refinement** |  |  |  |
| Initial model used (PDB code) | 7DXB | 7DXB | 7DXB |
| Model resolution (Å) | 2.9 | 2.9 | 2.9 |
| FSC threshold | 0.5 | 0.5 | 0.5 |
| Model resolution range (Å) | 250-2.9 | 250-2.9 | 250-2.9 |
| Map sharpening *B* factor (Å<sup>2</sup>) | -98.6 | -133.6 | -122.0 |
| Model composition |  |  |  |
| Non-hydrogen atoms | 23,064 | 23,312 | 23,168 |
| Protein residues | 2,884 | 2,896 | 3,136 |
| Ligands | 16 | 20 | 18 |
| *B* factors (Å<sup>2</sup>) |  |  |  |
| Protein | 86.19 | 136.02 | 208.96 |
| Ligand | 107.17 | 123.23 | 150.35 |
| R.m.s. deviations |  |  |  |
| Bond lengths (Å) | 0.003 | 0.004 | 0.006 |
| Bond angles (°) | 0.535 | 0.921 | 0.980 |
| Validation |  |  |  |
| MolProbity score | 1.47 | 1.46 | 1.65 |
| Clashscore | 8.68 | 8.58 | 7.84 |
| Poor rotamers (%) | 0.00 | 0.64 | 0.72 |
| Ramachandran plot |  |  |  |
| Favored (%) | 98.73 | 98.60 | 96.53 |
| Allowed (%) | 1.27 | 1.40 | 3.47 |
| Disallowed (%) | 0.00 | 0.00 | 0.00 |

